# Interfered Chromosome Pairing Promotes Meiosis Instability of Autotetraploid Arabidopsis by High Temperatures

**DOI:** 10.1101/2021.08.31.458414

**Authors:** Huiqi Fu, Jiayi Zhao, Ziming Ren, Ke Yang, Chong Wang, Xiaohong Zhang, Ibrahim Eid Elesawi, Xianhua Zhang, Jing Xia, Chunli Chen, Ping Lu, Yongxing Chen, Hong Liu, Guanghui Yu, Bing Liu

## Abstract

Alterations of environmental temperature affect multiple meiosis processes in flowering plants. Polyploid plants derived from whole genome duplication (WGD) have enhanced genetic plasticity and tolerance to environmental stress, but meanwhile face a challenge for organization and segregation of doubled chromosome sets. In this study, we investigated the impact of increased environmental temperature on male meiosis in autotetraploid *Arabidopsis thaliana*. Under low to mildly-increased temperatures (5-28°C), irregular chromosome segregation universally takes place in synthesized autotetraploid Columbia-0 (Col-0). Similar meiosis lesions occur in autotetraploid rice (*Oryza sativa* L.) and allotetraploid canola (*Brassica napus* cv. Westar), but not in evolutionary-derived hexaploid wheat (*Triticum aestivum*). As temperature increases to extremely high, chromosome separation and tetrad formation are severely disordered due to univalent formation caused by suppressed crossing-over. We found a strong correlation between tetravalent formation and successful chromosome pairing, both of which are negatively correlated with temperature elevation, suggesting that increased temperature interferes with crossing-over prominently by impacting homolog pairing. Besides, we showed that loading irregularities of axis proteins ASY1 and ASY4 co-localize on the chromosomes of *syn1* mutant, and the heat-stressed diploid and autotetraploid Col-0, revealing that heat stress affects lateral region of synaptonemal complex (SC) by impacting stability of axis. Moreover, we showed that chromosome axis and SC in autotetraploid Col-0 are more sensitive to increased temperature than that of diploid Arabidopsis. Taken together, our study provide evidence suggesting that WGD without evolutionary and/or natural adaption negatively affects stability and thermal tolerance of meiotic recombination in *Arabidopsis thaliana*.

## Introduction

Meiosis is a specialized type of cell division that, in plants, occurs in pollen mother cells (PMCs) and/or megasporocytes giving rise to gametes with a halved ploidy. At early stages of meiosis, meiotic recombination takes place between homologous chromosomes leading to exchange of genetic information via formation of crossovers (COs). Meiotic recombination results in novel combination of genetic alleles, which enables natural selection can happen in the progenies, and safeguards balanced chromosome segregation that is vital for production of viable gametes and fertility (Wang and Copenhaver, 2018). Meiotic recombination is initiated by generation of DNA double-strand breaks (DSBs) catalyzed by SPO11, that is a type-II topoisomerase (topoisomerase VI, subunit A) conserved among eukaryotes (Bergerat et al., 1997; Grelon et al., 2001; Stacey et al., 2006; Da Ines et al., 2020). Plants with defective DSB formation exhibit impaired homolog synapsis and recombination, and are male sterile due to mis-segregation of chromosomes (Grelon et al., 2001; Stacey et al., 2006; De Muyt et al., 2007; Xue et al., 2018; Da Ines et al., 2020). Subsequently, DSBs are repaired by recombinases RAD51 and DMC1, and eventually are processed into COs or non-COs (Klimyuk and Jones, 1997; Li et al., 2004; Sanchez-Moran et al., 2007; Pohl and Nickoloff, 2008; Da Ines et al., 2013; Singh et al., 2017; Su et al., 2017; Kobayashi et al., 2019; Yao et al., 2020). There are two types of COs, most of which belong to type-I class catalyzed by ZMM proteins (i.e., HEI10 and MLH1), and are spaced on chromosomes by interference (Higgins et al., 2004); the other COs (type-II) mediated by MUS81 are interference-insensitive (Hollingsworth and Brill, 2004; Berchowitz et al., 2007).

DSB formation and meiotic recombination rely on programmed building of chromosome axis and synaptonemal complex (SC). Meiotic-specific cohesion protein AtREC8/SYN1 is a key axis protein, dysfunction of which causes reduced DSB formation and impaired chromosome organization (Zickler and Kleckner, 1999; Lambing et al., 2020b). Two coiled-coil proteins ASY3 and ASY4 participate in organizing axis formation, and mediate the connections between the SYN1-mediated chromosome axis and SC through interplay with HORMA domain protein ASY1, which acts in DMC1-mediated meiotic recombination pathway (Armstrong et al., 2002; Sanchez-Moran et al., 2007; Ferdous et al., 2012; Chambon et al., 2018; Osman et al., 2018). ASY1 prevents preferential occurrence of COs at distal regions by antagonizing telomere-led recombination and maintaining CO interference (Lambing et al., 2020a). The conserved transverse filament protein ZYP1 composes the central element of SC and is crucial for homolog synapsis (Higgins et al., 2005; Wang et al., 2010; Barakate et al., 2014). Recent studies revealed that in Arabidopsis, ZYP1-dependent SC plays a role in shaping type-I CO rate by maintaining CO interference; moreover, the bias of CO rate between sexes is wiped when ZYP1 is knocked out (Capilla-Pérez et al., 2021; France et al., 2021).

Male meiosis in plants is sensitive to variations of environmental temperature (De Storme and Geelen, 2014; Bomblies et al., 2015; Liu et al., 2019; Lohani et al., 2019). In both dicots and monocots, low temperatures affect cytokinesis by disturbing the formation of phragmoplast, which thereby induces meiotic restitution and formation of unreduced gametes (Tang et al., 2011; De Storme et al., 2012; Liu et al., 2018). Under high temperatures, by comparison, both chromosome dynamics and cytokinesis are impacted; especially, meiotic recombination exhibits complex responses to thermal conditions (Draeger and Moore, 2017; Wang et al., 2017; Mai et al., 2019; De Storme and Geelen, 2020; Lei et al., 2020; Ning et al., 2021; Schindfessel et al., 2021). In Arabidopsis, a mild increase of temperature (28°C) positively affects type-I CO rate by enhancing the activity of ZMM proteins without impacting DSB formation (Lloyd et al., 2018; Modliszewski et al., 2018). At higher temperature (32°C), however, the rate and distribution of COs are altered and/or reshaped (De Storme and Geelen, 2020). When the temperature elevates beyond fertile threshold of Arabidopsis (36-38°C), CO formation is fully suppressed due to inhibited DSB generation and impaired homolog synapsis (Ning et al., 2021). Environmental temperatures therefore may manipulate genomic diversity, and/or influence stability of plant ploidy over generations by impacting male meiosis during microsporogenesis (Bomblies et al., 2015; Lohani et al., 2019).

Most higher plants, especially for angiosperms, have experienced at least one episode of whole genome duplication (WGD) event, which is considered an important force driving speciation, diversification, and domestication (Dubcovsky and Dvorak, 2007; Leitch and Leitch, 2008; Del Pozo and Ramirez-Parra, 2015; Soltis et al., 2015; Ren et al., 2018). Polyploids are classified into autopolyploids and allopolyploids, which originate from intraspecies WGD events, or arise from multiple evolutionary lineages through the combination of differentiated genomes, respectively (Bretagnolle and Thompson, 1995; Ramsey and Schemske, 1998; Soltis and Soltis, 2009; Jackson and Chen, 2010; Parisod et al., 2010). In autotetraploid plants, four intraspecies-homologues usually undergo randomly separation at anaphase I; this is different as allotetraploids, in which subgenomes tend to segregate independently due to preferential CO formation between the genetically-closer pairs of homologues (Ramsey and Schemske, 2002; Stift et al., 2008). It is considered that the duplicated genome contribute to genomic flexibility and confer plants with enhanced tolerance to both endogenous genetic mutations, or exogenous environmental stresses (Comai, 2005; te Beest et al., 2012; Del Pozo and Ramirez-Parra, 2015; Rao et al., 2020; Van de Peer et al., 2020; Wu et al., 2020). However, the multiple sets of homologous chromosomes also challenge genome stability by impacting chromosome pairing and segregation with associated reduced fertility or viability of plants (Santos et al., 2003; Comai, 2005; Otto, 2007; Yant et al., 2013; Svačina et al., 2020). It is proposed that evolutionarily-derived polyploids have developed a moderate strategy that assures genome stability to a large scale by early-stage homoeologous chromosome sorting, modification of chromosome axis-mediated meiotic recombination, and/or by sacrificing CO rate within an acceptable range (Grandont et al., 2014; Bomblies et al., 2016; Lloyd and Bomblies, 2016; Morgan et al., 2020; Seear et al., 2020). But, it remains not yet clear how male meiosis in polyploid plants behave s under thermal stress.

Here, we comprehensively analyzed meiosis behaviors of colchicine-induced autotetraploid Arabidopsis under increased temperatures. We found that a minor proportion of meiosis defects generally takes place in autotetraploid Col-0 under a wide range of temperature conditions, which are significantly increased when temperature reaches extremely high. We showed that heat stress interferes with meiosis in autotetraploid Arabidopsis via a prominent impact on chromosome pairing. Cytological analysis revealed that impaired homolog pairing and synapsis, and suppressed CO formation are owing to inhibited DSB generation and SC formation. Moreover, our data supported that heat stress destabilizes lateral structure of SC by targeting chromosome axis, and additionally suggested that stability of chromosome axis and SC in autotetraploid Arabidopsis is more sensitive to heat stress than that in diploid Arabidopsis. Overall, our findings propose that WGD without natural adaption negatively affects stability and thermal tolerance of meiotic recombination in *Arabidopsis thaliana*, which should be taken into consideration when applying polyploid breeding under the current fast climate-changing background.

## Results

### Heat stress increases aberrant meiotic products in autotetraploid *Arabidopsis thaliana*

To address the impact of heat stress on meiosis of autotetraploid Arabidopsis, we employed autotetraploid Col-0 plants generated by colchicine treatment on diploid Col-0 as previously reported (De Storme and Geelen, 2011). Fluorescence in situ hybridization (FISH) using a centromere-specific probe in somatic cells confirmed that the plants assayed were tetraploid (Supplemental Fig. S1). Under control temperature (20°C), most PMCs in autotetraploid Col-0 produced tetrads as diploid Col-0 (Fig. 1A-C). Interestingly, every single autotetraploid Col-0 plant was found to yield a small proportion of abnormal meiotic products (Fig. 1A, 4.39%; D and E), which was significantly increased under an extreme high temperature (37°C) (Fig. 1A, 93.91%; F-M). Aniline blue staining of tetrad stage meiocytes confirmed occurrence of defective meiotic cytokinesis in autotetraploid Col-0 under both temperature conditions (Supplemental Fig. S2).

**Figure 1.**
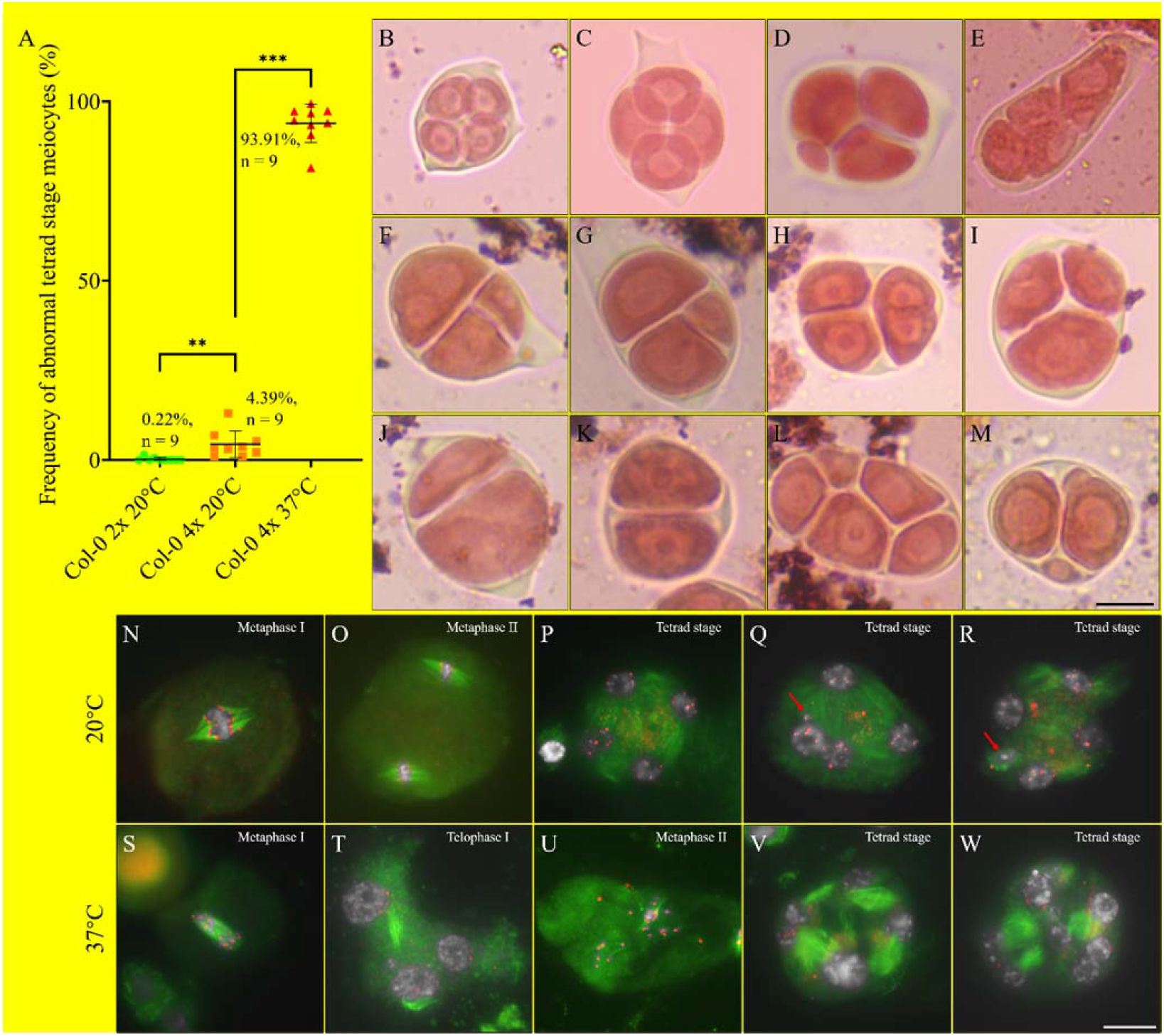
Analysis of PMCs in autotetraploid Col-0 plants. A, Graph showing the frequency of abnormal tetrad stage meiocytes in autotetraploid Col-0 plants incubated at 20°C and 37°C. The numbers indicate the frequency of abnormal tetrad PMCs; n indicates numbers of biological replicates used for quantification; ** and *** indicate *P* < 0.01 and 0.001, respectively. B-E, Orcein staining of tetrad stage meiocytes in diploid Col-0 (B) and autotetraploid Col-0 plants (C-E) grown at 20°C. F-M, Orcein staining of tetrad stage meiocytes in autotetraploid Col-0 plants stressed by 37°C. N-R, Immunolocalization of ɑ-tubulin in meiocytes at metaphase I (N), metaphase II (O) and tetrad (P-R) stages, respectively, of autotetraploid Col-0 plants incubated at 20°C. S-W, Immunolocalization of ɑ-tubulin in meiocytes at metaphase I (S), anaphase I (T), metaphase II (U) and tetrad (V and W) stages, respectively, of autotetraploid Col-0 plants stressed by 37°C. Red arrow indicates mini-nucleus. White, DAPI; green, ɑ-tubulin; red, CENH3. Scale bars = 10 μm.

Microtubular cytoskeleton was examined by performing immunolocalization of ɑ-tubulin. At 20°C, one and two sets of spindles were built at metaphase I and II, respectively, to separate homologs and sister chromatids (Fig. 1N and O). At telophase II, mini-phragmoplast structures composed of radial microtubule arrays (RMAs) were formed between the four isolated nuclei (Fig. 1P). Notably, as seen by orcein staining, triad- and polyad-like cells were observed, which showed omitted RMA between the two adjacent nuclei (and mini-nucleus, red arrow), or irregular RMA formation between the multiple nuclei (Fig. 1Q and R). After heat treatment, assembly of spindles at metaphase I and II was interfered (Fig. 1S and U); meanwhile, phragmoplast at anaphase I displayed aberrant orientation and sparse microtubule content (Fig. 1T). Most tetrad stage meiocytes exhibited impaired RMA formation (Fig. 1V and W). These findings suggested that male meiosis is unstable in the synthesized autotetraploid Arabidopsis grown under normal temperature conditions, which, additionally, is hypersensitive to heat stress.

### Heat-induced meiosis defects are highly correlated with interfered chromosome pairing

Since Arabidopsis plants defective for meiotic recombination (e.g., the *dmc1* mutant) also show disorganized spindle and phragmoplast during meiosis (Supplemental Fig. S3) (Bai et al., 1999; Xue et al., 2019; Shi et al., 2021), heat-induced abnormalities in autotetraploid Col-0 may result from alterations in meiotic recombination. In naturally-derived autotetraploid Arabidopsis (*A. arenosa*), meiotic recombination rate varies in response to seasonal temperature changes (Weitz et al., 2021). Driven by the curiosity how meiotic chromosomes in synthesized autotetraploid *Arabidopsis thaliana* would behave under increased temperature, we applied meiotic spreading analysis in autotetraploid Col-0 shocked by a wide range of low-to-high temperatures (i.e., 5, 10, 15, 28, 32 and 37°C, respectively). First, pachytene stage chromosomes were examined to see the impact of temperature elevation on homolog pairing. We found that pairing defect (i.e., homologous chromosomes were not fully paired) existed in autotetraploid Col-0 grown at any given temperature conditions (Fig. 2A-C). Bridge- and/or thick rope-like structures implied an improper chromosome pairing (Fig. 2D and E). The frequency of pairing lesions did not show any significant difference at lower temperatures (i.e., 5-20°C) (Fig. 2A). However, as temperature increased, pairing abnormalities were significantly pronounced (Fig. 2A and F), and there was no pairing at 37°C (Fig. 2A, *P* < 0.01; G). The rate of successful pairing showed a strong negative correlation with elevated temperatures (Fig. 3A).

**Figure 2.**
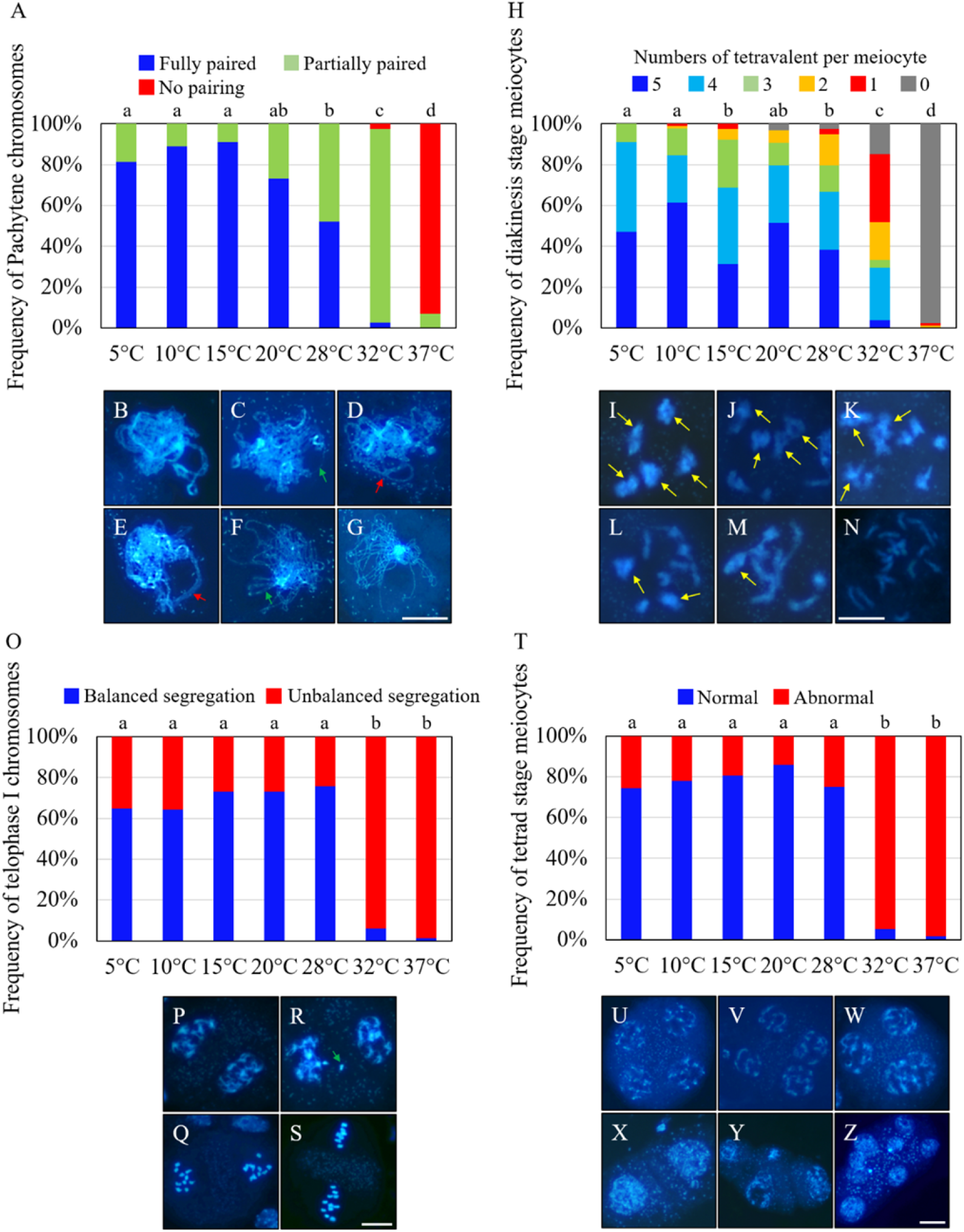
Meiotic chromosome behaviors in autotetraploid Col-0 shocked by different temperatures. A, Graph showing the frequency of fully-paired, partially-paired and no-pairing pachytene chromosomes in autotetraploid Col-0 at different temperatures. B-G, Pachytene stage chromosomes showing fully-paired (B), partially-paired (C), irregularly-associated (D and E), minorly-paired (F) and non-paired (G) configurations, respectively. Green arrows indicate unpaired chromosomes; red arrows indicate irregularly-associated chromosomes. H, Graph showing the frequency of diakinesis stage meiocytes with varied tetravalent formation in autotetraploid Col-0 at different temperatures. I-N, Diakinesis stage meiocytes with five (I), four (J), three (K), two (L), one (M) and no (N) tetravalents, respectively. Yellow arrows indicate tetravalents. O, Graph showing the frequency of telophase I stage meiocytes showing balanced and/or unbalanced homolog segregation in autotetraploid Col-0 at different temperatures. P-S, Interkinesis (P and R) and metaphase II (Q and S) stage meiocytes showing balanced (P and Q) and unbalanced (R and S) chromosome segregation, respectively. Green arrow indicates lagged chromosome. T, Graph showing the frequency of normal and/or abnormal tetrad stage meiocytes in autotetraploid Col-0 at different temperatures. U-Z, Tetrad stage meiocytes showing normal (U) and abnormal (V-Z) configurations. Nonparametric and unpaired *t* test was performed. Lower letters indicate significant difference between different temperatures by setting significance level *P* < 0.05. Original quantification data and number of cells quantified were supplied in Supplement Table S1-S4. Scale bars = 10 μm.

**Figure 3.**
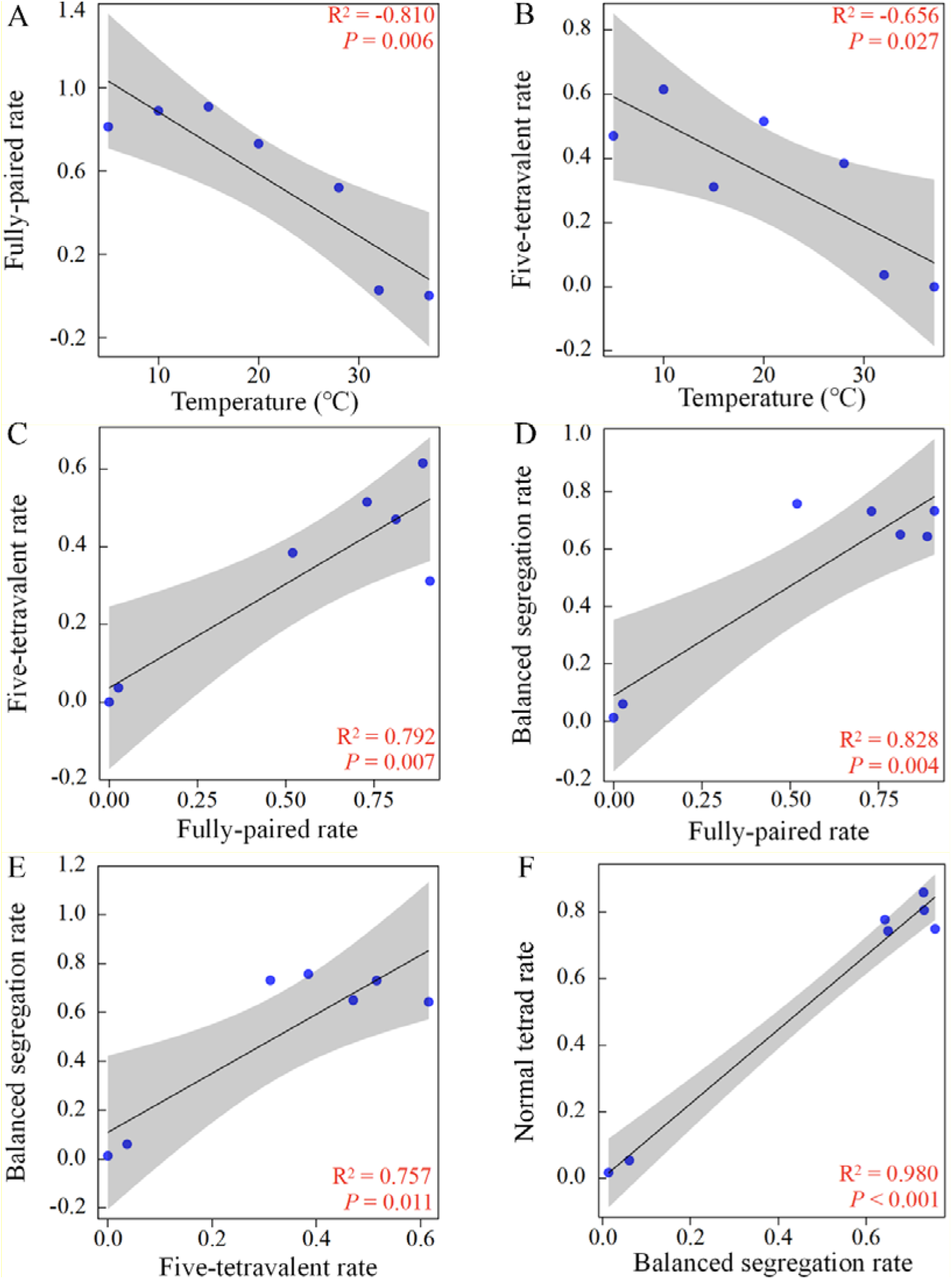
Correlation between chromosome behaviors of autotetraploid Col-0 at different temperature conditions. A and B, Correlations between frequency of chromosome pairing (A) and five-tetravalent formation (B) with temperature variations. C and D, Correlations between frequency of five-tetravalent formation (C) and balanced chromosome segregation at anaphase I (D) with rate of fully-pairing of pachytene chromosomes. E, Correlation between frequency of balanced chromosome segregation at anaphase I with rate of five-tetravalent formation. F, Correlation between frequency of normal tetrad formation with rate of balanced chromosome segregation at anaphase I.

At diakinesis stage, autotetraploid Col-0 plants under low-to-normal temperatures (i.e., 5-20°C) preferentially generate PMCs with five tetravalents, the rate of which was decreased when the temperature climbed to extremely high (Fig. 2H, *P* < 0.01; I-N; Fig. 3B). A positive correlation was found between rate of five-tetravalent formation and successful chromosome pairing (Fig. 3C). Under all the temperatures tested, unbalanced chromosome segregation occurred after meiosis I and II, and showed a significant high level under extreme thermal conditions (Fig. 2O and T, *P* < 0.01; P-S, U-Z). The similar rate of heat-induced irregularities at telophase I and tetrad stage suggested that heat stress induces abnormal meiotic products predominantly by impairing chromosome segregation at meiosis I (Fig. 2O and T; Fig. 3F). Balanced chromosome separation was positively correlated with chromosome pairing status and the ratio of five-tetravalent formation (Fig. 3D and E). Taken together, these data thus suggested that heat stress interferes with chromosome segregation in autotetraploid *Arabidopsis thaliana* via a primary impact on chromosome pairing.

To address whether meiosis lesions in synthesized autotetraploid Arabidopsis generally occurs in polyploid plants, we checked chromosome behaviors in meiocytes of colchicine-induced autotetraploid rice, naturally-derived allotetraploid canola and allohexaploid wheat grown at cultivation temperatures. In both autotetraploid rice and allotetraploid canola, irregular chromosome separation at anaphase I and/or tetrad stage were visualized (Supplemental Fig. S4F; Supplemental Fig. S5A-H); nevertheless, no obvious meiosis defect was observed in allohexaploid wheat (Supplemental Fig. S6A-J). Interestingly, in autotetraploid rice, pachytene stage chromosomes did not show pairing defects (Supplemental Fig. S4A and B), but displayed irregularly arranged chromosomes at metaphase I cell plate (Supplemental Fig. S4C and D, red arrow). These findings hence imply that meiosis instability universally takes place in polyploid plants without evolutionary adaption and/or with genetic instability (Lu et al., 2019).

### Type-I CO rate is lowered in autotetraploid Arabidopsis under high temperatures

Univalent formation in autotetraploid Col-0 plants under high temperatures suggested that CO formation was compromised or suppressed. To this end, we performed immunolocalization of HEI10 protein, which marks class I-type CO (Chelysheva et al., 2012; Wang et al., 2012), and quantified its abundance on the diakinesis chromosomes. At 20°C, autotetraploid Col-0 showed an average of 18.10 HEI10 foci per meiocyte (Fig. 4A and B), which was reduced to an average of 11.80 and 0.56 in the plants stressed by 32°C and 37°C, respectively (Fig. 4C and D). This data indicated that high temperatures inhibit formation of class I-type CO in autotetraploid Arabidopsis.

**Figure 4.**
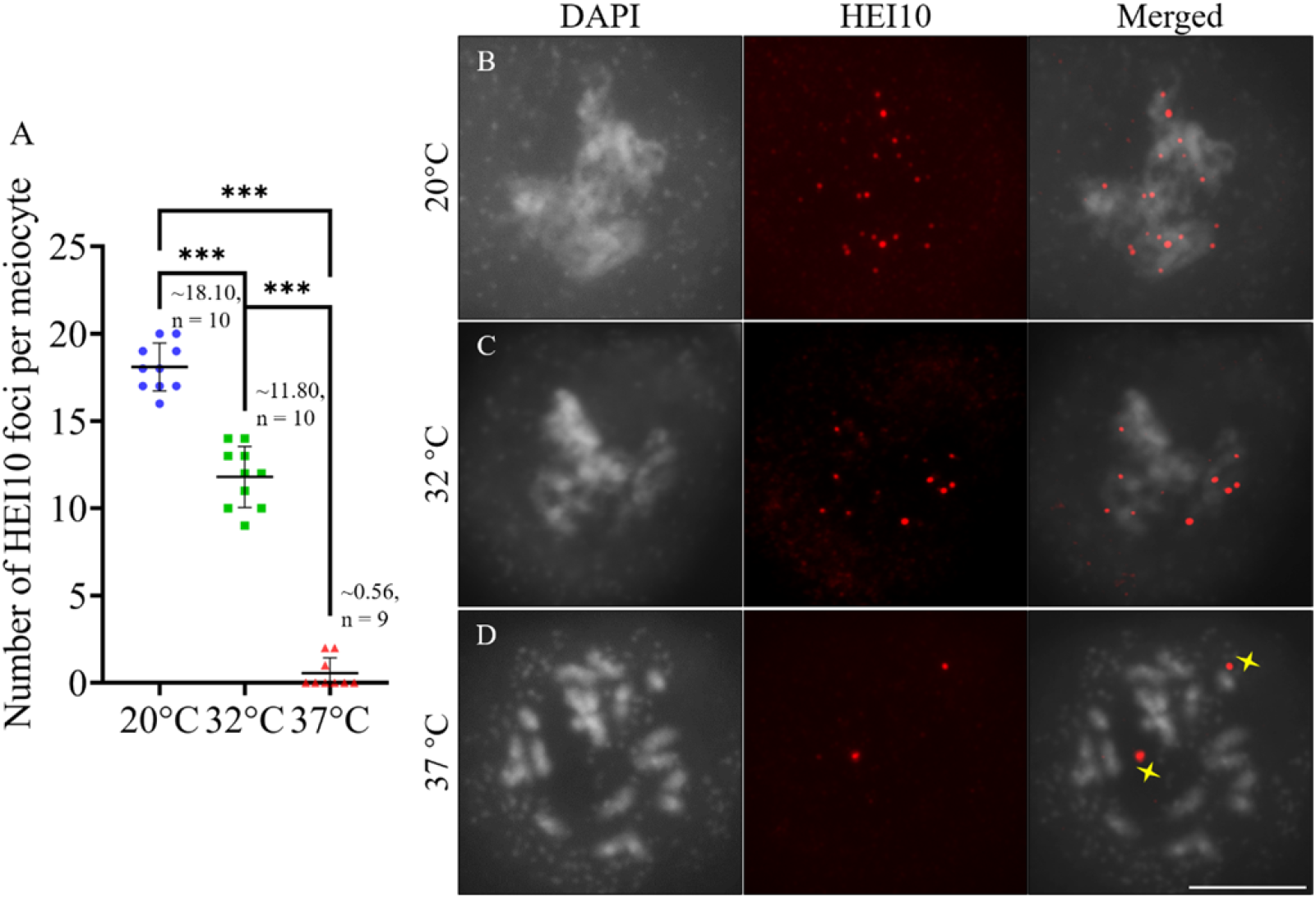
Immunolocalization of HEI10 on diakinesis chromosomes in autotetraploid Col-0 plants under increased temperatures. A, Graph showing the number of HEI10 foci per diakinesis-staged meiocyte. Numbers indicate the average number of HEI10 foci per meiocyte; n indicates the number of cells quantified; *** indicates *P* < 0.001. B-D, Immunolocalization of HEI10 on diakinesis chromosomes of autotetraploid Col-0 plants incubated at 20°C (B), 32°C (C) and 37°C (D). Yellow stars indicate non-specific foci to HEI10. Scale bar = 10 μm.

### Heat stress impairs SC assembly in autotetraploid Arabidopsis

SC assembly is required for homolog synapsis (Higgins et al., 2005; Wang et al., 2010;Barakate et al., 2014; Capilla-Pérez et al., 2021; France et al., 2021), we subsequently examined central element of SC in heat-stressed autotetraploid Col-0 plants by performing immunolocalization of ZYP1 protein, the core element of transverse filament. At 20°C,zygotene meiocytes displayed partially-assembled linear ZYP1 configuration (Fig. 5A). Thereafter ZYP1 were fully assembled at the central region of the paired homologs at middle pachytene, and were gradually disassociated as disintegration of SC from late pachytene (Fig. 5B and C). By contrast, plants stressed by 37°C displayed dotted and/or fragmented installation of ZYP1 from early zygotene to late pachytene (Fig. 5D-H, yellow arrow), which indicated that assembly of SC was impaired. Aggregated and/or enlarged ZYP1 foci hinted pairing between multiple chromosomes (Fig. 5D, G and H, blue arrows) (Loidl,1989;Morgan et al., 2017;Ning et al., 2021). Localization of SYN1 did not show any obvious defect indicating that heat stress does not affect SYN1-mediated axis (Fig. 5D-H). Moreover, in line with the chromosome spreading analysis, ZYP1 loading was not influenced by cold (5°C) (Supplemental Fig. S7A-C), but was slightly impacted under mildly increased temperature (28°C) (i.e., incomplete ZYP1 loading occurring at unpaired chromosome regions (Supplemental Fig. S7D-F, red arrow).

**Figure 5.**
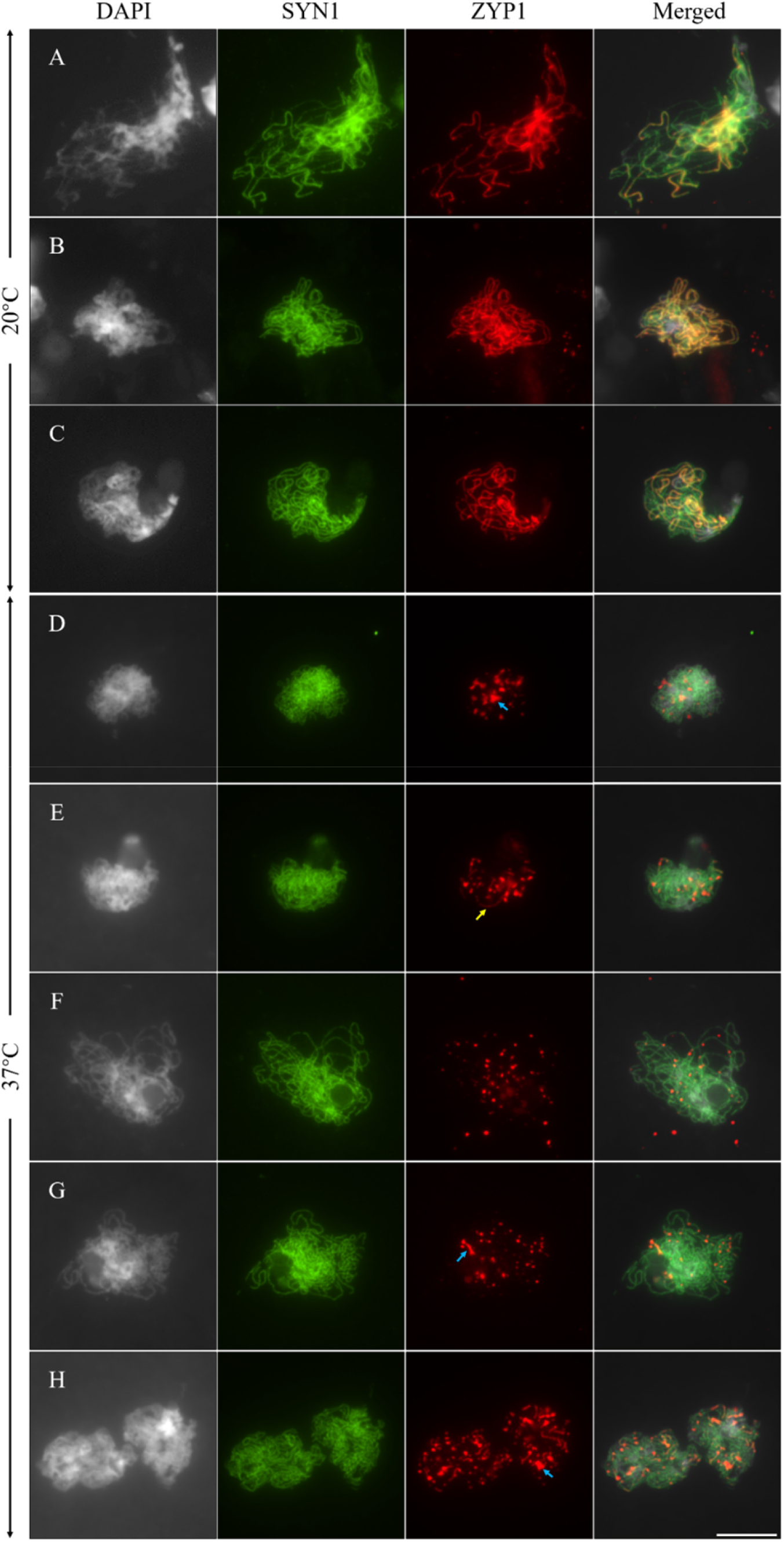
Co-immunolocalization of SYN1 and ZYP1 in meiocytes of autotetraploid Col-0 plants. A-C, Zygotene- (A), middle pachytene- (B) and late pachytene-staged (C) meiocytes in autotetraploid Col-0 at 20°C. D-H, Early zygotene- (D), middle zygotene- (E), late zygotene- (F), middle pachytene- (G) and late pachytene-staged (H) meiocytes in autotetraploid Col-0 stressed by 37°C. Green, SYN1; red, ZYP1. Yellow arrow indicates fragmented ZYP1 signals. Blue arrows indicate aggregated and/or enlarged ZYP1 foci. Scale bar = 10 μm.

### Heat stress reduces DSB formation in autotetraploid Arabidopsis

DSB generation is crucial for homolog pairing and CO formation (De Muyt et al., 2007; Hartung et al., 2007; Kurzbauer et al., 2012). To address whether heat-interfered chromosome pairing and CO reduction was owing to compromised DSB formation, we indirectly scored DSB abundance by counting the number of ɤH2A.X, which specifically marks phosphorylated histone variant H2A.X at DSB sites (Kurzbauer et al., 2012). Autotetraploid Col-0 incubated at 20°C showed an average of 146.5 ɤH2A.X foci per meiocyte at zygotene, and was lowered to 79.46 and 84.8 per meiocyte when the temperature was elevated to 32°C and 37°C, respectively (Fig. 6A-D). In addition, the number of DMC1 on zygotene chromosomes decreased under the high temperatures, either (Fig. 6E-H). These data indicated that DSB formation is compromised in heat-stressed autotetraploid Arabidopsis.

**Figure 6.**
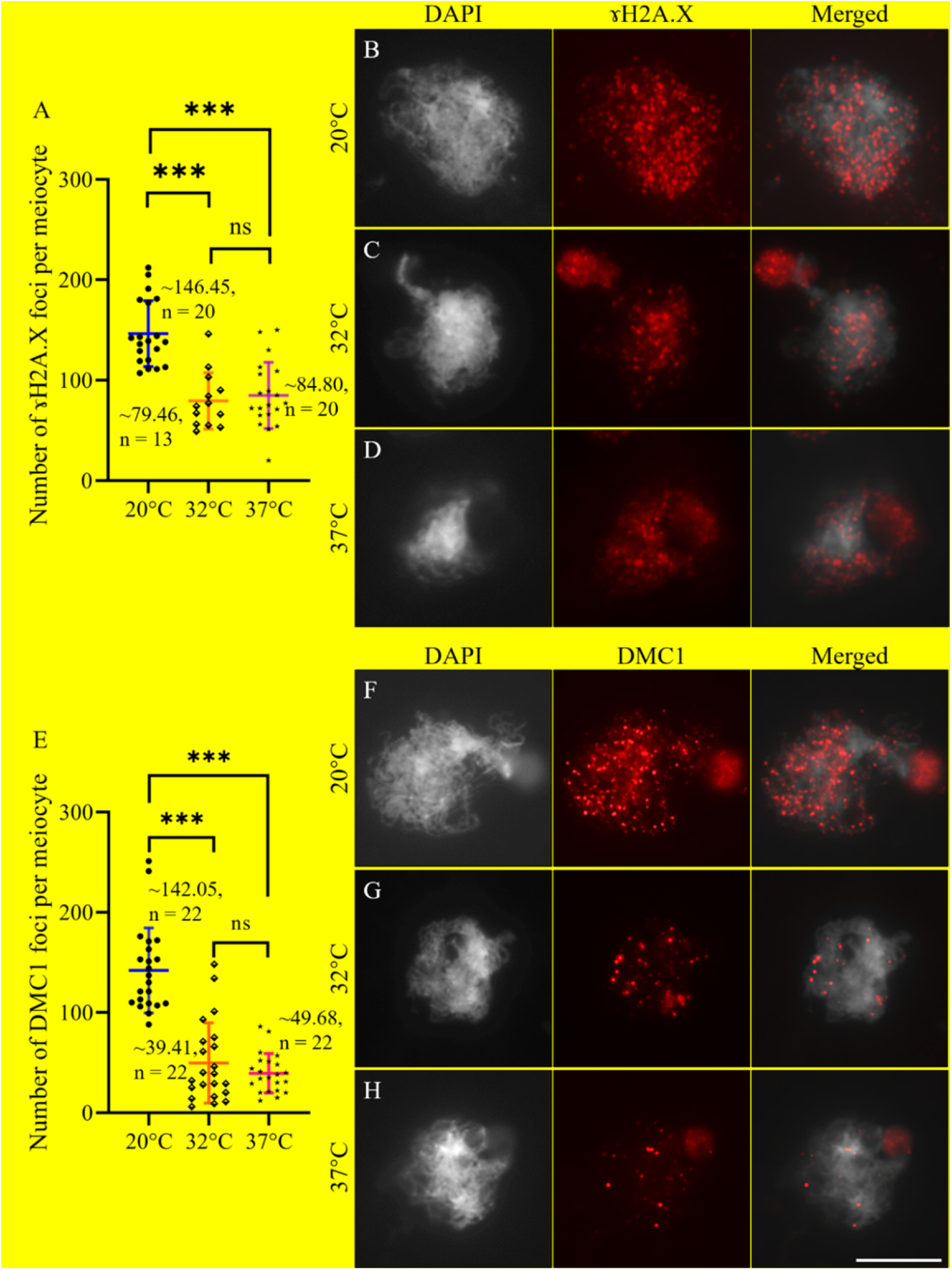
Immunolocalization of ɤH2A.X and DMC1 on zygotene chromosomes of autotetraploid Col-0 plants. A, Graph showing the number of ɤH2A.X foci per meiocyte in autotetraploid Col-0 plants at 20°C, 32°C and 37°C. B-D, Immunolocalization of ɤH2A.X on zygotene chromosomes of autotetraploid Col-0 plants grown at 20°C (B), 32°C (C) and 37°C (D), respectively. E, Graph showing the number of DMC1 foci per meiocyte in autotetraploid Col-0 plants at 20°C, 32°C and 37°C. F-H, Immunolocalization of DMC1 on zygotene chromosomes of autotetraploid Col-0 plants at 20°C (F), 32°C (G) and 37°C (H), respectively. Numbers indicate the average number of ɤH2A.X or DMC1 foci per meiocyte; n indicates the number of cells quantified; *** indicates *P* < 0.001; ns indicates no significant difference. Scale bars = 10 μm.

### Higher heat-sensitivity of chromosome axis in autotetraploid Arabidopsis

Successful homolog synapsis and CO formation also rely on normal assembly of chromosome axis. Since SYN1 is not impacted by heat stress, we examined the dynamics of two other axis-associated proteins (i.e., ASY1 and ASY4) in autotetraploid Col-0. In control plants, linear SYN1 and ASY1 overlapped and were associated with the entire chromosomes at zygotene (Fig. 7A and B). ASY1 were unloaded at some chromosome regions at pachytene when homolog fully synapsed (Fig. 7C and D). After heat treatment, most zygotene and pachytene chromosomes in autotetraploid Col-0 displayed dotted configuration of ASY1, whose frequency was significantly higher than that in heat-stressed diploid Col-0 (Fig. 7E-H; Supplemental Fig. S8; Supplemental Fig. S10A-C). Similar phenotypic alterations have been observed in ASY4 loading in heat-stressed autotetraploid Col-0 (Fig. 8A-E; Supplemental Fig. S9; Supplemental Fig. S10D-F). These hinted that ASY1- and ASY4-mediated chromosome axis in autotetraploid Arabidopsis is more sensitive to heat than that in diploid Arabidopsis.

**Figure 7.**
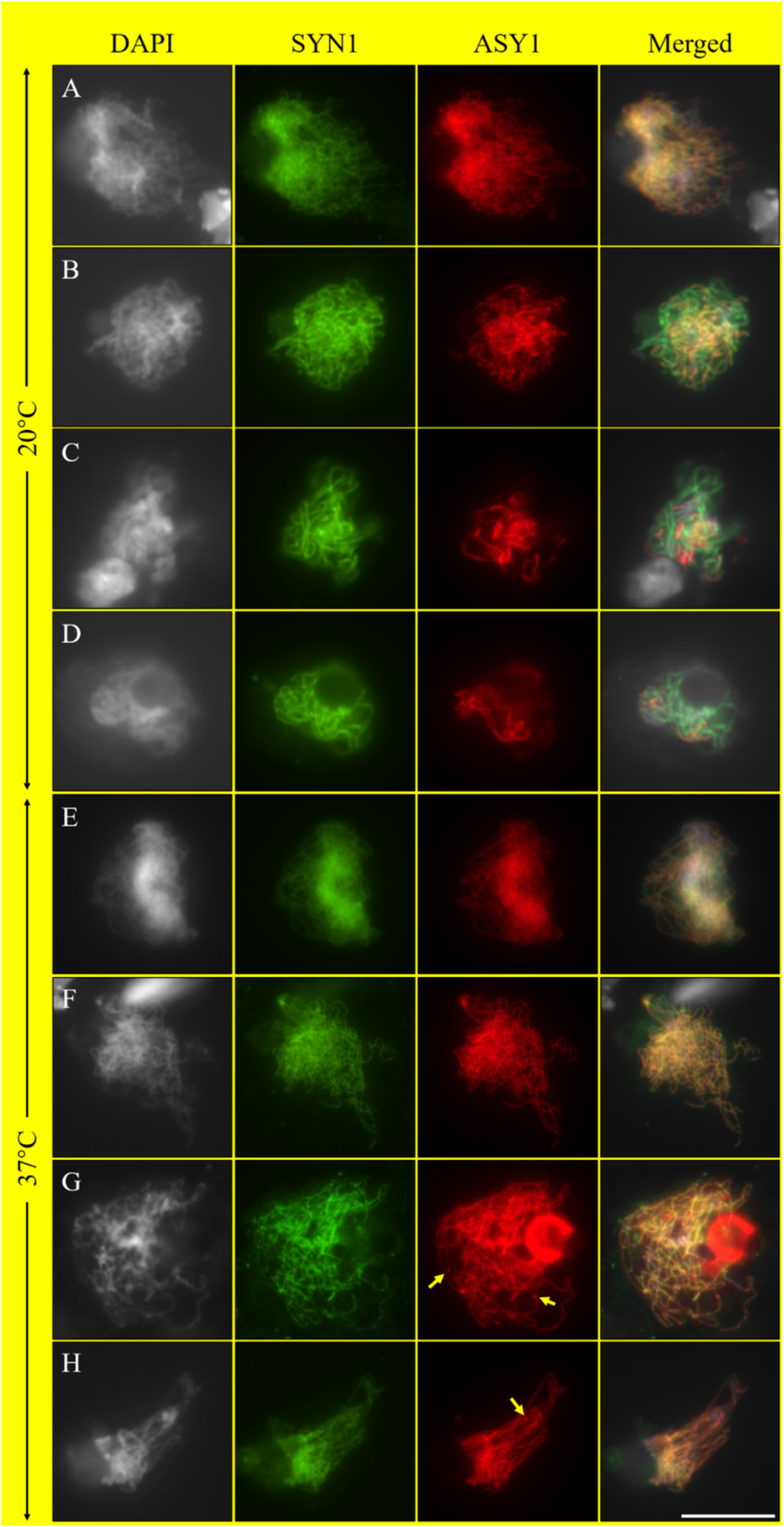
Co-immunolocalization of SYN1 and ASY1 in meiocytes of autotetraploid Col-0 plants. A-D, Early zygotene (A), middle zygotene (B), middle pachytene (C) and late pachytene (D) chromosomes in autotetraploid Col-0 plants at 20°C. E-H, Zygotene (E and G) and pachytene (F and H) chromosomes showing linear (E and F) and dotted (G and H) configurations, respectively, in autotetraploid Col-0 plants at 37°C. Green, SYN1; red, ASY1. Yellow arrows indicate dotted ASY1 foci. Scale bar = 10 μm.

**Figure 8.**
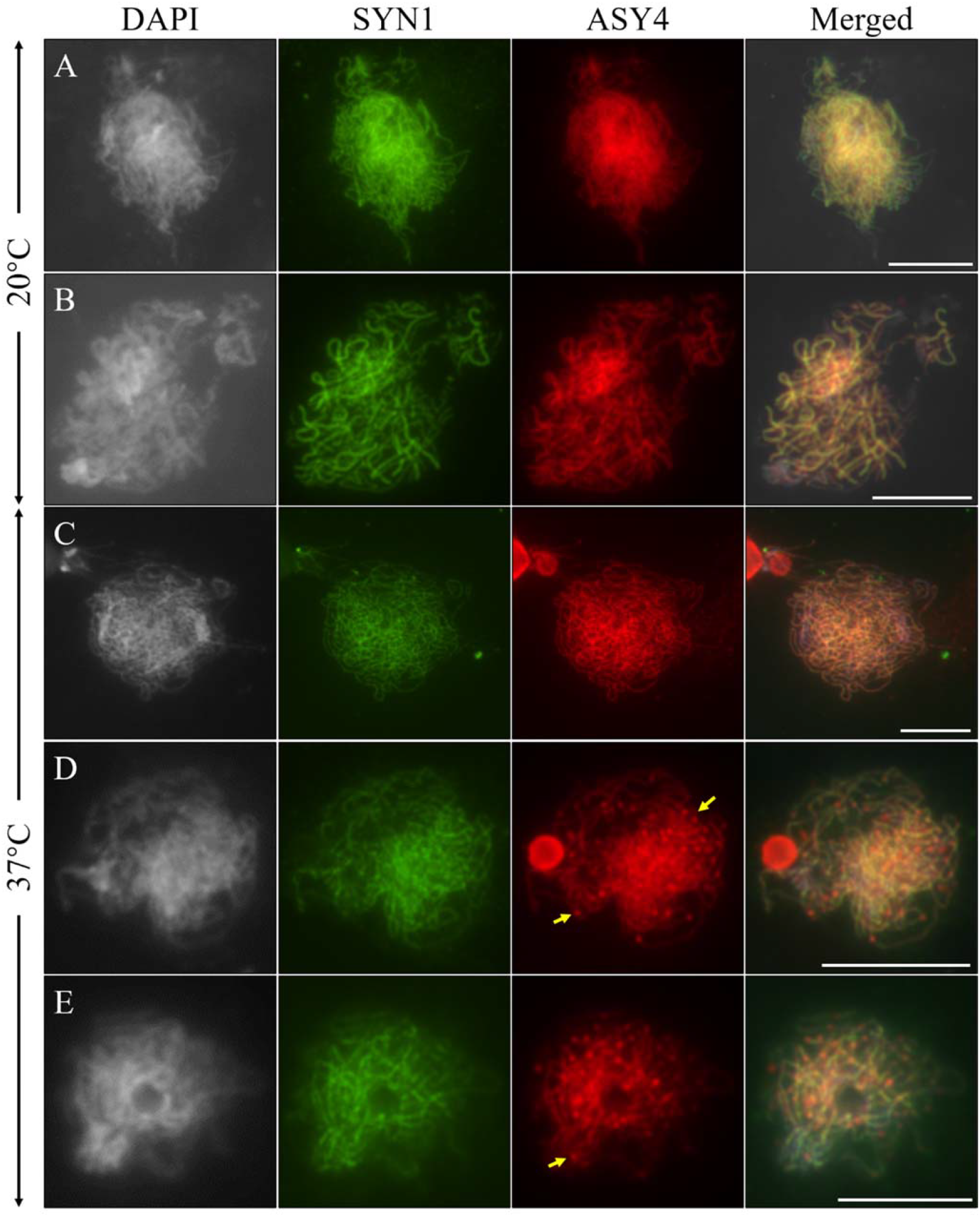
Co-immunolocalization of SYN1 and ASY4 in meiocytes of autotetraploid Col-0 plants. A and B, Zygotene (A) and pachytene (B) chromosomes in autotetraploid Col-0 plants at 20°C. C-E, Zygotene (C and D) and pachytene (E) chromosomes showing linear (C) and dotted (D and E) configurations, respectively, in autotetraploid Col-0 plants at 37°C. Green, SYN1; red, ASY4. Yellow arrows indicate dotted ASY4 foci. Scale bars = 10 μm.

### Heat stress affects lateral element of SC by impacting stability of chromosome axis

It is proposed that assembly of ASY1-associated lateral element of SC relies on a step-wise formation of SYN1-ASY3-ASY4-mediated chromosome axis (Ferdous et al., 2012; Chambon et al., 2018; Lambing et al., 2020b). We performed co-immunolocalization of ASY1 and ASY4 in diploid Col-0, and the *syn1, spo11-1-1, rad51* and *dmc1* mutants (Fig. 9). In the wild-type and the *spo11-1-1, rad51* and *dmc1* mutants, ASY1 and ASY4 co-localize on middle zygotene chromosomes (Fig. 9A; D-F). By contrast, the *syn1* mutant displayed incomplete and/or fragmented ASY1 and ASY4 loading, which additionally overlapped on the zygotene and pachytene chromosomes (Fig. 9B and C). These observations are in line with the opinion that DSB processing is downstream of axis assembly, and ASY1-associated SC formation relies on ASY4-mediated axis formation, which in turn depends on functional SYN1.

**Figure 9.**
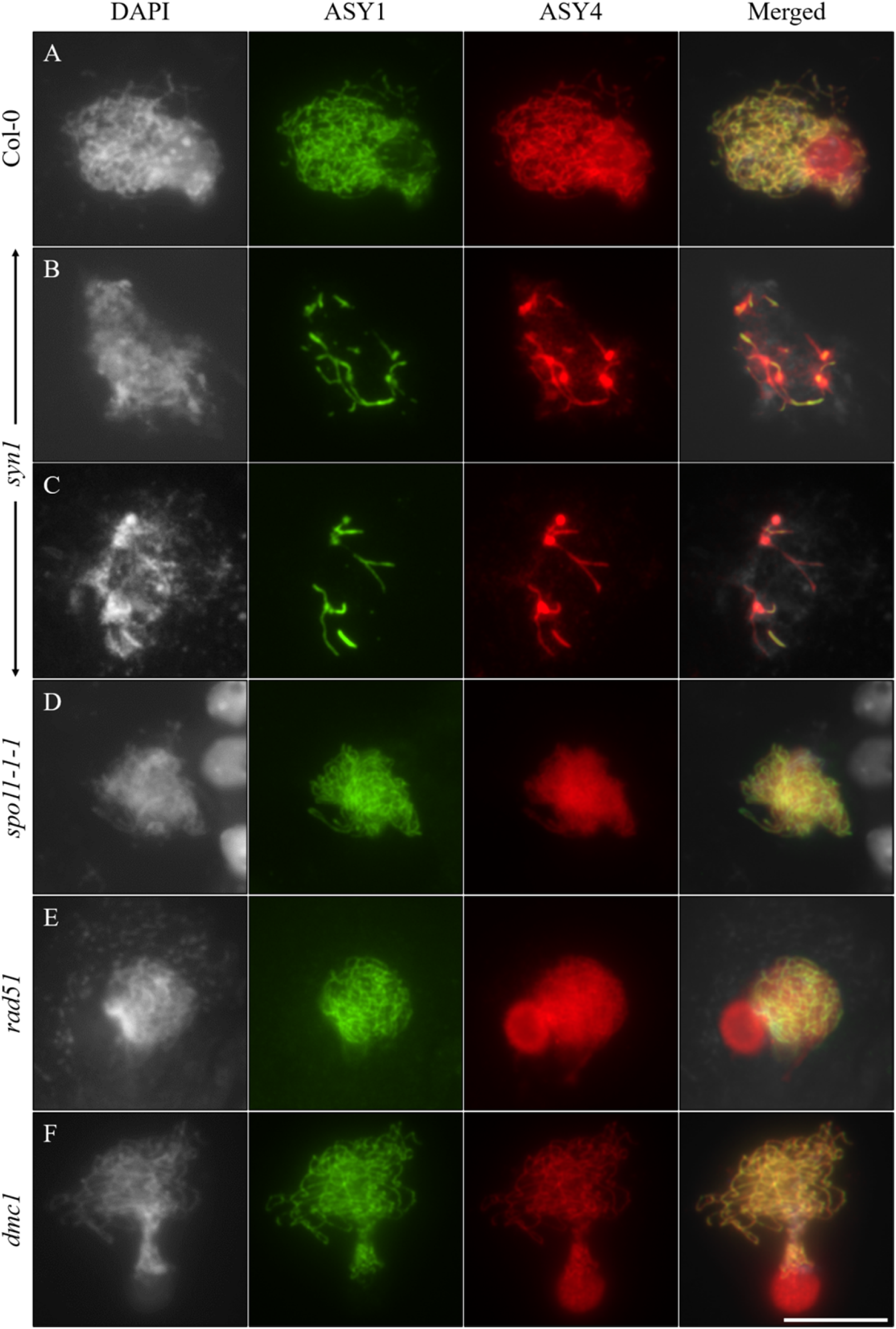
Co-immunolocalization of ASY1 and ASY4 in meiocytes of diploid wild-type Col-0, and the *syn1, spo11-1-1, rad51* and *dmc1* mutants. A-F, Zygotene chromosomes in diploid Col-0 (A); zygotene (B) and pachytene (C) chromosomes in the *syn1* mutant; zygotene chromosomes in the *spo11-1-1* (D), *rad51* (E) and the *dmc1* (F) mutants. Green, ASY1; red, ASY4. Scale bar = 10 μm.

Considering the similarities of defective ASY1 and ASY4 loading under heat stress, and the upstream action of axis formation on SC assembly, we hypothesized that heat stress destabilizes ASY1-associated lateral regions of SC via an impacted ASY4 stability. To this end, a combined immunostaining of ASY1 and ASY4 was applied in the heat-stressed diploid and autotetraploid Col-0 plants. Under control temperature, ASY1 and ASY4 co-localized on the early and middle zygotene chromosomes of diploid and autotetraploid Col-0 (Fig. 10A, diploid Col-0; D, autotetraploid Col-0; Supplemental Fig. S11A and B, autotetraploid Col-0). Subsequently, ASY1 started to be unloaded off the chromosomes from early pachytene, while ASY4 kept a complete linear configuration (Supplemental Fig. S11C-E), indicating that ASY4 is disassembled later than ASY1 (Supplemental Fig. S11F and G). Notably, in both the diploid and autotetraploid Col-0, heat-induced incomplete and/or dotted ASY1 and ASY4 signals co-localized on the chromosomes (Fig. 10B and C, diploid Col-0; E and F, autotetraploid Col-0; yellow arrows). It thus is likely that heat stress affects ASY1-associated SC via compromised stability of ASY4-mediated chromosome axis.

**Figure 10.**
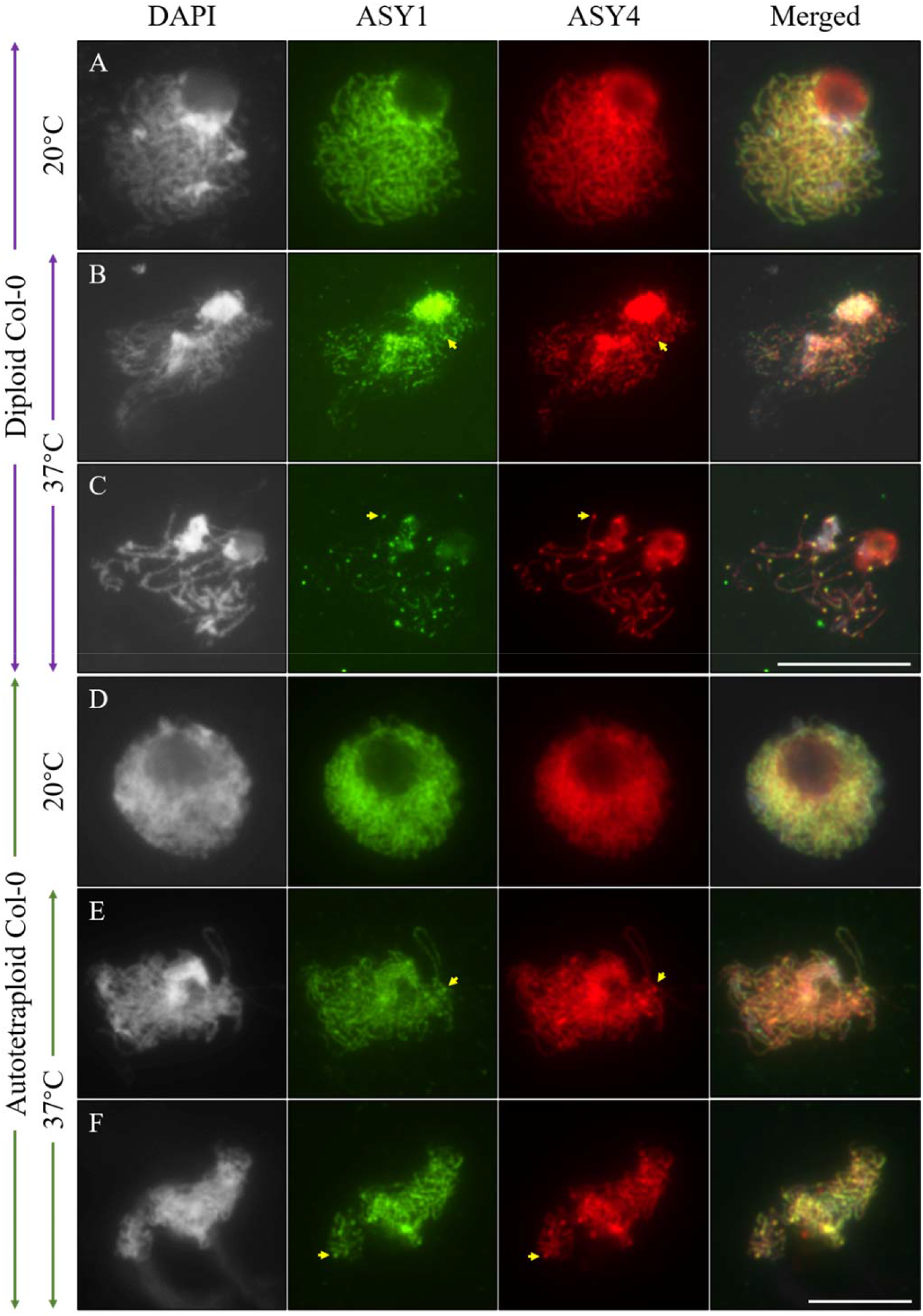
Co-immunolocalization of ASY1 and ASY4 in heat-stressed diploid and autotetraploid Col-0 plants. A and D, Zygotene-staged meiocytes in diploid (A) and autotetraploid (D) Col-0 plants grown at 20°C. B and C, Zygotene-(B) and pachytene-staged (C) meiocytes in heat-stressed diploid Col-0. E and F, Zygotene- (E) and pachytene-staged (F) meiocytes in autotetraploid Col-0 at 37°C. Green, ASY1; red, ASY4. Yellow arrows indicate co-localization of dotted ASY1 and ASY4 foci. Scale bars = 10 μm.

## Discussion

### Meiosis is unstable in autotetraploid *Arabidopsis thaliana* without natural adaption

WGD is a conserved phenomenon that contributes to genomic diversity and speciation in higher plants (Comai, 2005; Dubcovsky and Dvorak, 2007; te Beest et al., 2012; Ren et al., 2018; Van de Peer et al., 2020; Li et al., 2021). Additional copies of chromosomes, however, also increase the complexity and challenge for homolog pairing and synapse which eventually harms balanced chromosome segregation at later meiosis stages (Yant et al., 2013; Lloyd and Bomblies, 2016; Svačina et al., 2020; Li et al., 2021). In this study, we found that synthesized autotetraploid Col-0 yields a low but consistently-detectable rate of aberrant meiotic products under normal temperature conditions (Fig. 1). Chromosome analysis revealed that unsuccessful pairing and unbalanced chromosome segregation occur in autotetraploid Arabidopsis grown under a wide range of low-to-high temperatures (Fig. 2). The co-aligned, but un-synapsed axis regions suggest that the synapsis defects are (or at least in part) independent of defects in chromosome pairing (Fig. 2F) (Capilla-Pérez et al., 2021; France et al., 2021). The autotetraploid Col-0 plants that we used here is very typical of neo-autotetraploids that are considered genetically unstable compared with the evolution-derived autotetraploids (e.g., *A. lyrate* and *A. arenosa*) (Yant et al., 2013; Henry et al., 2014; Lloyd and Bomblies, 2016). In support, synthesized autotetraploid rice and naturally-derived allotetraploid canola, which, however, is genetically unstable (Lu et al., 2019), also show defective segregation of homologous chromosomes (Supplemental Fig. S4F; Supplemental Fig. S5B-D). By contrast, meiosis in evolutionarily-derived hexaploid *Triticum aestivum* behaves normally (Supplemental Fig. S6) (El Baidouri et al., 2017). The meiotic defects we observed here hence are probably the effect of polyploidization without natural adaption and/or evolutionary selection. On the other hand, in synthesized autotetraploid rice, we did not find pachytene chromosomes with pairing defects (Supplemental Fig. S4A and B), suggesting that meiosis alterations in autotetraploid rice may not be caused by interfered homolog pairing but by other disorders; e.g., improper multivalent dissolution (Supplemental Fig. S4D) (Lloyd and Bomblies, 2016). Therefore, the mechanisms of WGD interfering with meiosis may vary among species.

### Increased temperatures impose a dominant impact on chromosome pairing

We found that the rate of successful chromosome pairing and tetravalent formation were not altered under 5-20°C, suggesting that homolog pairing and CO formation are more sensitive to high temperatures (Fig. 2A and H; Supplemental Fig. S7A-C). Autotetraploid Arabidopsis primarily generates diakinesis PMCs that contain five tetravalents (Fig. 2H), which is in line with the notion that autotetraploids preferentially undergo synapsis between four homologous chromosomes when environmental temperature is suitable for meiotic recombination (Lloyd and Bomblies, 2016; Svačina et al., 2020; Braz et al., 2021). The negative correlation between chromosome pairing, tetravalent formation and increased temperatures imply that high temperatures affect CO formation predominantly by interfering with homolog pairing (Fig. 3A-C). Similarly, in naturally-derived autotetraploid Arabidopsis, multivalent formation is strongly correlated with seasonal temperature alterations (Weitz et al., 2021). Polyploid plants thus may apply same (at least very similar) mechanism in response to climate changes during meiotic recombination.

Heat-induced DSB reduction could be one of the main causes of impaired chromosome pairing in heat-stressed Arabidopsis (Fig. 6), which, additionally, may be a conserved phenomenon among eukaryotes (Pohl and Nickoloff, 2008; Ning et al., 2021). But this effect is not likely to be caused by a direct impact on the expression of DSB formation factors (i.e., *SPO11-1, PRD1, 2* and *3*) (Ning et al., 2021). Notably, the expression of phosphatidylinositol 3 kinase-like (PI3K) protein kinase *Ataxia-Telangiectasia Mutated* (*ATM*), which undertakes a conserved function in sensing DNA damage and evoking DSB repair events (reviewed by (Paull, 2015)), is elevated under heat stress (Ning et al., 2021). In multiple species, the activity of ATM is negatively correlated with DSB abundance (Joyce et al., 2011; Lange et al., 2011; Zhang et al., 2011; Carballo et al., 2013; Garcia et al., 2015; Mohibullah and Keeney, 2017; Shi et al., 2019). In Arabidopsis, ATM limits DSB formation by restricting the accumulation of SPO11 on prophase I chromosomes (Yao et al., 2020; Kurzbauer et al., 2021). Therefore, it is possible that high temperatures interfere with DSB formation via an over-activated ATM function. On the other hand, our data suggest that high temperatures can directly disorder SC formation, since autotetraploid Arabidopsis stressed by 32°C has similar level of DSB as at 37°C, but exhibits a mildly-compromised rate of successful chromosome pairing and CO formation (Fig. 2, 4 and 6) (De Storme and Geelen, 2020).

### Heat stress affects lateral element of SC by impacting ASY4-mediated chromosome axis

The loading of ASY1 and ASY4 is compromised in the *syn1* mutant (Fig. 9B and C) (Ning et al., 2021), which, however, is not the case conversely (Ferdous et al., 2012; Chambon et al., 2018; Lambing et al., 2020b). Meanwhile, linear configuration of ASY3 depends on functional ASY4 (Chambon et al., 2018). Therefore, SYN1, ASY4 and ASY3 may act by a stepwise manner in mediating axis formation. However, since ASY4 directly interacts with ASY3, the normal loading of ASY4 thus may also rely on the existence of ASY3 (Chambon et al., 2018). The slower unloading of ASY4 than ASY1 on later prophase I chromosomes supports that SC assembly is downstream of axis formation (Supplemental Fig. S9) (Chelysheva et al., 2005; Ferdous et al., 2012; Chambon et al., 2018; Lambing et al., 2020a; Lambing et al., 2020b). Heat-induced ASY1 and ASY4 abnormalities occur at a similar frequency, which, meanwhile, co-localize on the chromosomes (Fig. 10; Supplemental Fig. S10). These data hence support the hypothesis that high temperatures destabilize lateral structure of SC via impacted chromosome axis. Since SYN1 keeps stable under the high temperatures (Fig. 5, 7 and 8) (Ning et al., 2021), it is plausible that heat stress specifically targets the ‘bridge’ function of ASY4 and/or ASY3 that associate chromosome axis with SC. Whether the impacted axis stability also channels the compromised DSB formation under heat stress remain further investigation.

### WGD does not increase thermal tolerance of meiosis in *Arabidopsis thaliana*

Polyploid plants are considered to have evolutionarily developed enhanced tolerance to genetic variations due to higher gene copies, and to abiotic stresses via modulated hormone metabolism and/or reprofiled gene expression (Allario et al., 2013; Lourkisti et al., 2020; Rao et al., 2020; Van de Peer et al., 2020). However, the significantly reduced abundance of γH2A.X and DMC1 in autotetraploid Col-0 plants under high temperatures suggest that the duplicated genome does not change the thermal threshold of DSB formation in *Arabidopsis thaliana* (Fig. 6) (Ning et al., 2021). The slightly increased defect of chromosome pairing at 28°C (Fig. 2A; Supplemental Fig. S7), which does not occur in diploid Arabidopsis (Modliszewski et al., 2018), implies that homolog pairing and/or synapsis in autotetraploid Arabidopsis is also more heat-sensitive. Meanwhile, autotetraploid Arabidopsis shows higher rate of impacted ASY1- and ASY4-mediated chromosome axis to heat shock (Supplemental Fig. S10). Our findings thus suggest that genome duplication does not promote thermal tolerance of meiotic recombination in *Arabidopsis thaliana*. Indeed, chromosome axis components plays an important role in maintaining meiosis stability in tetraploid Arabidopsis, whose expression is high temperature-sensitive (Morgan et al., 2020; Ning et al., 2021). Therefore, polyploidization-induced higher stress tolerance may not be a general event. One of the explanation could be that environmental stimulus modulates gene expression in a tissue-specific manner in polyploids (Adams and Wendel, 2005). Alternatively, enhanced axis instability under heat shock may be attributed to an increased complexity of chromosome organization in autotetraploid Arabidopsis plants which has not experienced evolutionary adaption (Morgan et al., 2020; Seear et al., 2020; Svačina et al., 2020).

## Materials and methods

### Plant materials and growth conditions

Diploid and autotetraploid *Arabidopsis thaliana* Columbia-0 (Col-0), autotetraploid rice (*O. sativa* L. ssp. *Indica* cv. 9311) (Gan et al., 2021), hexaploid *Triticum aestivum* cultivar ‘Fielder’, allotetraploid canola (*Brassica napus* cv. Westar), the *atsyn1-1* (SALK_137095), *atrad51* (SAIL_873_C08), *atspo11-1-1* (Grelon et al., 2001) and the *atdmc1* (SALK_056177) mutants were used in the study. The autotetraploid Col-0 plants were generated by colchicine treatment on diploid Col-0 plants as described previously (De Storme and Geelen, 2011), and have been propagated around five generations. Determination of chromosome number was performed in somatic cells by fluorescence in situ hybridization. Arabidopsis plants were grown under a 16 h day/8 h night, 20°C, and 50% humidity condition. Rice plants were grown in paddy fields during the growing season in Wuhan (30.52°N, 114.31°E), China. Canola and wheat were grown in a growth chamber with a 16 h day/8 h night and 22°C condition.

### Temperature treatment

Young flowering plants were transferred from control temperature (20°C) to a humid chamber with a 16 h day/8 h night and incubated at 5°C, 10°C, 15°C, 28°C, 32°C and 37°C, respectively, and were treated for 24 h. All the treatment started from 8:00-10:00 AM. Meiosis-staged flower buds were fixed by carnoy’s fixative or paraformaldehyde upon the finish of treatment.

### Cytology and fluorescence in situ hybridization

Meiotic chromosome behaviors were analyzed by performing chromosome spreading using meiosis-staged flower buds fixed at least 24 h by carnoy’s fixative. Flower buds were washed twice by distilled water and once in citrate buffer (10 mM, pH = 4.5), and were incubated in digestion enzyme mixture (0.3% pectolyase, 0.3% cellulase and 0.3% cytohelicase) in citrate buffer (10 mM, pH = 4.5) at 37°C in a moisture chamber for 2.5-3.5 h. Subsequently, 6-8 digested buds were washed in distilled water, and were transferred to a glass slide followed by squashing with a small amount (4-5 μL) of distilled water. Two rounds of 10 μL precooled 60% acetic acid were added to the samples and were stirred gently, after which the slide was transferred to a hotplate at 45°C for 1-2 min, and thereafter was flooded with precooled carnoy’s fixative. The slides were subsequently air dried for 10 min, and were stained by adding 8 μL DAPI (10 μg/mL) in Vectashield antifade mounting medium, mounted with a coverslip, and sealed by nail polish. To analyze and quantify meiotic products, tetrad-staged flower buds were stained by 45% orcein, and the flower buds containing significant number of tetrad-staged PMCs were used for quantification. Five biological replicates were analyzed both for control and heat-stressed plants. To analyze meiotic cytokinesis, tetrad-staged flower buds were stained by aniline blue (0.1% in 0.033% K_3_PO_4_). FISH assay and the centromere-specific probe have previously been reported (Lei et al., 2020). For rice, canola and wheat, young panicles were fixed with carnoy’s fixative, and anthers containing PMCs occurring meiosis were squashed followed by 45% orcein staining or chromosome spreading analysis.

### Generation of antibodies

The anti-AtSYN1 antibodies were raised in rabbits and mouse, respectively, as previously reported (Bai et al., 1999); the anti-AtASY1 antibodies were generated in rabbits and mouse, respectively, against amino acid sequence SKAGNTPISNKAQPAASRES of AtASY1 conjugated to KLH; the anti-AtZYP1 antibody (rat) was generated against the amino acid sequence GSKRSEHIRVRSDNDNVQD of AtZYP1A conjugated to KLH.

### Immunolocalization of MR proteins and ɑ-tubulin

Immunostaining of ɑ-tubulin and MR proteins was performed as reported (Chelysheva et al., 2010; ang et al., 2014; Liu et al., 2017). Antibodies against ZYP1 (rabbit and/or rat) (Ning et al., 2021), HEI10 (rabbit), DMC1 (rabbit) (Ning et al., 2021) and γH2A.X (rabbit) (Lambing et al., 2020b) were diluted by 1:100; antibodies against ɑ-tubulin (rat) (Lei et al., 2020), ASY1 (rabbit and/or mouse), ASY4 (rabbit) (Ning et al., 2021) and SYN1 (mouse) were diluted by 1:200; antibody against CENH3 (rabbit) (Abcam, 72001) was diluted by 1:400; antibody against SYN1 (rabbit) was diluted by 1:500. The secondary antibodies; i.e. Goat anti-Rabbit IgG (H+L) Cross-Adsorbed Secondary Antibody Alexa Fluor 555 (Invitrogen, A32732), Goat anti-Rabbit IgG (H+L) Highly Cross-Adsorbed Secondary Antibody Alexa Fluor Plus 488 (Invitrogen, A32731), Goat anti-Rat IgG (H+L) Cross-Adsorbed Secondary Antibody, Alexa Fluor 555 (Invitrogen, A21434), Goat anti-Rat IgG (H+L) Cross-Adsorbed Secondary Antibody, Alexa Fluor 488 (Invitrogen, A11006) and Goat anti-Mouse IgG (H+L) Highly Cross-Adsorbed Secondary Antibody, Alexa Fluor Plus 488 (Invitrogen, A32723) were diluted to 10 µg/mL.

### Microscopy and quantification of fluorescent foci

Bright-field images and DAPI-stained meiotic chromosomes were pictured using a M-Shot ML31 microscope equipped with a MS60 camera. Aniline blue staining of meiotic cell walls, and immunolocalization of ɑ-tubulin and MR-related proteins were analyzed on an Olympus IX83 inverted fluorescence microscope equipped with a X-Cite lamp and a Prime BSI camera. Image processing and quantification of fluorescent foci were conducted as previously reported (Ning et al., 2021). Briefly, images of DAPI-stained chromosome signals and RFP-channeled protein foci signals were merged, and only the foci merged onto chromosomes were considered the specific foci to targeted proteins, and were counted manually using the Image J count tool.

### Statistical analysis

To analyze the significant difference of orcein-stained tetrad stage meiocytes, meiotic spreading phenotypes, ɤH2A.X, DMC1 and HEI10 foci counts, and ASY1/ASY4 configurations under different temperatures, nonparametric and unpaired *t* test was performed using the software GraphPad Prism (version 8). Correlation analysis was conducted using SPSS (IBM, 22.0, the U.S.). Pearson’s correlation was used to determine correlation coefficients. * and ** represent *P* value < 0.05 and 0.01, respectively. Figures were produced using the R statistical platform (version 3.6.3) via R studio software (version 1.3.1093), and package ggplot2 (version 3.3.3, Wickham, 2009) was used for visualization. The number of cells and/or the biological replicates used for quantification were indicated in the figures and/or supplemental materials.

## Supplemental materials

Supplemental Figure S1. FISH analysis of somatic cells in autotetraploid Col-0.

Supplemental Figure S2. Meiotic cell wall formation in autotetraploid Col-0 stressed by 37°C.

Supplemental Figure S3. Immunolocalization of ɑ-tubulin in the *dmc1* mutant.

Supplemental Figure S4. Meiotic spreading analysis of meiocytes in autotetraploid rice.

Supplemental Figure S5. Meiotic spreading analysis of meiocytes in allotetraploid canola.

Supplemental Figure S6. Orcein-staining of PMCs in hexaploid *Triticum aestivum*.

Supplemental Figure S7. Immunolocalization of ASY1 and ZYP1 in meiocytes of autotetraploid Col-0 at 5 and 28°C.

Supplemental Figure S8. Immunolocalization of ASY1 in meiocytes of autotetraploid Col-0 at 37°C.

Supplemental Figure S9. Immunolocalization of ASY4 in meiocytes of autotetraploid Col-0 at 20°C.

Supplemental Figure S10. Quantification of ASY1 and ASY4 configurations in diploid and autotetraploid Col-0 stressed by 37°C.

Supplemental Figure S11. Immunolocalization of ASY1 and ASY4 in meiocytes of autotetraploid Col-0 at 20°C.

Supplemental Table S1. Quantification of pachytene stage meiocytes in autotetraploid Col-0.

Supplemental Table S2. Quantification of diakinesis stage meiocytes in autotetraploid Col-0.

Supplemental Table S3. Quantification of anaphase I stage meiocytes in autotetraploid Col-0.

Supplemental Table S4. Quantification of tetrad stage meiocytes in autotetraploid Col-0.

Supplemental Table S5. Primers used in the study.

## Acknowledgement

The authors thank Dr. Jing Li (Huazhong Agricultural University), Dr. Detian Cai (Hubei University) and Dr. Chao Yang (Huazhong Agricultural University) for kindly providing the autotetraploid Col-0 seeds, autotetraploid rice and allotetraploid canola samples, respectively. They thank Dr. Yingxiang Wang (Fudan University) for kindly sharing the anti-HEI10 (rabbit) antibody. They appreciate Dr. Andrew Lloyd (IBERS) for critical review and comments on the manuscript prior to submission.

## Author contribution

H.Q.F. and J.Y.Z. performed immunolocalization experiments; Z.M.R. performed statistical analysis and contributed to data interpretation; K.Y. and X.H.Z. contributed to chromosome spreading analysis; C.W. analyzed fluorescent foci; E.I.E. performed FISH experiment; X.H.Z., J.X., C.L.C., P.L. Y.X.C., H.L. and G.H.Y. contributed to data analysis; B.L.conceived and designed the study, analyzed data, wrote, and edited the manuscript.

## Funding

This work was supported by National Natural Science Foundation of China (32000245 to B.L.), Hubei Provincial Natural Science Foundation of China (2020CFB159 to B.L.), Fundamental Research Funds for the Central Universities, South-Central University for Nationalities (CZY20001 to B.L.), Fundamental Research Funds for the Central Universities, South-Central University for Nationalities (YZZ18007 to B.L.), National Natural Science Foundation of China (31900261 to C.W.), National Natural Science Foundation of China (31270361 to G.H.Y.), Fundamental Research Funds for the Central Universities (CZZ21004 to G.H.Y.), and National Natural Science Foundation of China (31971525 to C.L.C.).

## Interest of conflict

All the authors declared that there is no conflict of interest in this work.

